# High-volume, label-free imaging for quantifying single-cell dynamics in induced pluripotent stem cell colonies

**DOI:** 10.1101/2023.09.29.558451

**Authors:** Anthony Asmar, Zack Benson, Adele P. Peskin, Joe Chalfoun, Mylene Simon, Michael Halter, Anne Plant

## Abstract

To facilitate the characterization of unlabeled induced pluripotent stem cells (iPSCs) during culture and expansion, we developed an AI pipeline for nuclear segmentation and mitosis detection from phase contrast images of individual cells within iPSC colonies. The analysis uses a 2D convolutional neural network (U-Net) plus a 3D U-Net applied on time lapse images to detect and segment nuclei, mitotic events, and daughter nuclei to enable tracking of large numbers of individual cells over long times in culture. The analysis uses fluorescence data to train models for segmenting nuclei in phase contrast images. The use of classical image processing routines to segment fluorescent nuclei precludes the need for manual annotation. We optimize and evaluate the accuracy of automated annotation to assure the reliability of the training. The model is generalizable in that it performs well on different datasets with an average F1 score of 0.94, on cells at different densities, and on cells from different pluripotent cell lines. The method allows us to assess, in a non-invasive manner, rates of mitosis and cell division which serve as indicators of cell state and cell health. We assess these parameters in up to hundreds of thousands of cells in culture for over 20 hours, at different locations in the colonies, and as a function of excitation light exposure.

## Introduction

Induced pluripotent stem cells (iPSC) can be created from any adult cell, they can expand almost indefinitely, and they can be induced to differentiate into any cell type of the body. iPSC therapies are being developed to treat diseases, reconstruct tissues, and screen drugs (1). The culture, expansion, and differentiation of iPSCs in either a research or manufacturing setting is challenging because of their sensitivity to conditions such as media composition, physical handling (such as passaging protocols), and the variations that exist between different cell lines. This presents the need for better quantification, and real-time monitoring, of the effects of culture conditions on stem cell renewal, expansion, and maintenance of pluripotency. Time-lapse imaging of live cells is a powerful quantitative tool for longitudinal measurements of iPSCs.

Fluorescence labeling is a powerful tool in cell imaging and in the creation of image analysis algorithms, but exposure of cells for long times to potentially harmful excitation irradiation is undesirable (2–6). The non-destructive nature of low-intensity transmitted light imaging allows for real-time, relatively non-invasive, optical probing of living cells in culture over long times, but thus far it has not been possible to segment and track individual IPSC by transmitted light imaging alone. Furthermore, if adequate analytical pipelines can be established and validated using low-intensity transmitted light, any cell line can be quantitatively imaged over time in an non-invasive manner. Metrics such as morphology, migration rate, division times, and expression of reporter molecules could be used for characterizing and monitoring the state of cells in real-time during culture and expansion processes. Optical imaging that provides such metrics would allow the testing and development of unique culture and expansion conditions that are optimal for different cell lines. The spatial and temporal nature of live cell imaging provides contextual knowledge that is not available from single timepoint and non-imaging measurements.

The growth of pluripotent cells in multicellular colonies has posed challenges to quantitative imaging at the cellular level because of their close cell-cell contacts. Imaging at the colony level (7) or as groups of cells (8) in brightfield and fluorescence have led to knowledge about stem cell colony growth characteristics, gene expression, and classification at different stages of stem cell culture. Neural network-based algorithms have enabled greater discrimination of the characteristics of groups of iPSC and have been successfully used to predict differentiation (9) or distinguish between iPSC colonies, differentiated cells and dead cells from phase contrast images in an automated expansion culture setting (10–12). In another study, an assessment of individual iPSC nuclei based on a fluorescent label and an ensemble of neural networks addressed the challenge of dense cell tracking and localization of each cell nucleus in an iPSC colony but required the signal of a fluorescent probe (13). We present here quantitative dynamic imaging and tracking of unlabeled individual iPSCs and their progeny over time.

We previously developed a neural network-based workflow for segmentation of single iPSC nuclei from label-free, transmitted light microscopy images (14). The training data did not require manual annotation but rather utilized a reporter cell line expressing a green fluorescent protein (GFP)-nuclear protein fusion product (LaminB-GFP) to enable nuclear segmentation. This approach for automated segmentation allowed generation of large training data sets and an evaluation of model performance as a function of the amount of training data. Inferenced nuclear detection incurred false positive rates and false negative rates of 0.16 and 0.19, respectively. More recently we developed a 3D neural network mitosis detection model for dividing iPS cells from label-free, transmitted light microscopy images (15). The model generation depended on manual annotations and a unique semi-supervised approach to iteratively expand the training data size and improve model performance. The final model successfully identified 37 of the 38 cell division events in manually annotated test image stack.

In this study, we build on our prior work to report a label-free, transmitted light acquisition and analysis workflow for segmenting and tracking individual iPSC nuclei, including mitosis detection for characterizing the onset of mitosis, division, and parent-daughter cell linkages. This workflow required refinement of our previous models to improve performance, and the use of a cell line expressing a different fluorescent nuclear probe (H2BJ-mEGFP), which enabled improved mitosis detection for annotating phase contrast images for training the models. Both training and evaluation of the models rely on large datasets of auto-segmented images of fluorescent nuclei.

Ultimately, we envision using this analysis pipeline to enable the label-free tracking and quantification of reporters of gene expression in individual cells over time. Because the complexity and stochastic nature of cell regulatory pathways results in heterogeneous responses from different cells, it is critical to collect data on statistically sufficient numbers of individual cells, and this requirement precludes heavily supervised image analysis. Here our goal is to minimize the use of human intervention while efficiently collecting large sets of data for training and evaluating the segmentation and tracking of label-free iPS cells.

## Methods

### Cell culture

A genetically engineered cell line expressing green fluorescent protein (mEGFP) covalently fused to the histone protein, HIST1H2BJ, was developed by Allen Institute for Cell Science (mEGFP-tagged Histone H2B type 1-j) and procured from Coriell (AICS-0061-036, Philadelphia). Parental iPSCs (WTC-11, GM25256, Coriell) and hESCs (WA09, WiCell, Madison WI) were maintained under feeder-free conditions on Matrigel-coated plates (#354277, BD Biosciences) in mTeSR™ Plus (#100-0276, StemCell Technologies Inc.). iPSCs were routinely passaged with Accutase (# 07920, StemCell Technologies Inc.), while hESCs were routinely passaged with collagenase (#07909, StemCell Technologies Inc.). For experimental setups, cells were passaged as single cells using Accutase, enumerated using a Multisizer 3 Coulter Counter (Beckman,), and then seeded at the desired density in 6-well tissue culture plates (TPP, Trasadingen, Switzerland). U-Net models were trained using data with cells seeded at 5,200 cells/cm^2^(low density), 10,400 cells/cm^2^(medium density), and 20,800 cells/cm^2^ (high-density). All live cell experiments were performed on cells seeded at approximately 10,400 cells/cm^2^. Spy595-DNA (CY-SC301; Cytoskeleton, Denver CO) was used to label nuclei for generating reference data for WTC-11 and H9 cells according to manufacturer’s instructions.

### Image acquisition workflow

Zernike phase and fluorescence images were collected using a 10X/0.3 numerical aperture objective (Zeiss part number 420341-9911-000) on a Zeiss Axio Observer.Z1 microscope (Carl Zeiss USA, Thornwood, NY) equipped with motorized x-y stage (SCANplus IM 130x80, Marzhauser, Germany) and an ORCA-Fusion BT Digital CMOS camera (C15440-20UP, Hamamatsu, Japan) for image capture. Images were typically collected at 2min intervals. For each time point, multiple images are acquired (2000x2000 pixels per field of view) with 10% overlap between tiles. Each field-of view contains 7 phase contrast images at different z-planes and a single fluorescence image. A spatial calibration target was used to determine that each pixel is equivalent to an area of 0.406 µm^2^. During automated acquisition, the microscope and all peripherals were controlled with an electronics and software controller system that enabled faster data acquisition than is typically achieved with instrumentation software provided by most manufacturers (Inscoper, Rennes, France). A detailed description of the acquisition protocols and benchmarking of the microscope system is provided in the Supplemental Material. A description of experimental datasets including the experiment name, light exposure, cell counts, number of mitoses, and length of experiment, see Supplemental Table 1.

### Generating nuclear object reference data

Typical datasets include hundreds of images in a time-lapse acquisition, where each stitched, single timepoint image contains tens of thousands of cells. This volume of data presents a sampling challenge for assessing the performance of a label-free nuclear segmentation algorithm. Manually annotating thousands of cells would yield less than a 0.1% sampling rate. To address this big data challenge, we used classical segmentation routines to automate the creation of reference objects from GFP fluorescence images for testing the performance of our U-Net models.

The reference masks for training the 2D U-Net are generated by segmenting the GFP fluorescence image using a band-pass filter followed by an empirically determined intensity threshold value. To separate touching objects, the mask objects are segmented with a watershed-like segmentation routine, FogBank (16) followed by a binary erosion.

### Training the 2D U-Net

We trained a 2D U-Net to segment single-cell nuclei from phase contrast images starting with a pre-trained U-Net (14) as our initial network. Training data consisted of a 15000x15000 pixel subset of a large-scale single timepoint stitched phase contrast image (approximately 230 million pixels), and a corresponding fluorescence image where nuclei are labeled with H2B-GFP.

The phase contrast images are normalized using z-score normalization, and the final images and reference masks are tiled into 256x256 tiles to match the input of the U-Net. The relative weights of the foreground and background classes is 2 to 1.

Image augmentation was used during the training process in which the input training data were modified by operations such as reflection, rotations, and by applying Gaussian blur to mimic potential variations in cell orientation and focus. Training of the U-Net was done in MATLAB (https://github.com/usnistgov/WSDOM).

### Inferencing for label-free segmentation

The trained U-Nets operate on normalized phase contrast images and return binary foreground/background masks corresponding to single-cell nuclei. The binary masks are created by inferencing with 3 instances of the same model and thresholded by 2 (as explained in more detail in Supplemental Figure 1Band 1C.) The binary masks are post-processed using FogBank segmentation (16) to separate touching objects. Fogbank includes two input parameters, the erosion size and the minimum size. The effect of these parameters on nuclear detection performance was evaluated (Supplemental Figure 1A). We chose parameters that maximized the performance for the rest of this study.

**Figure 1.**
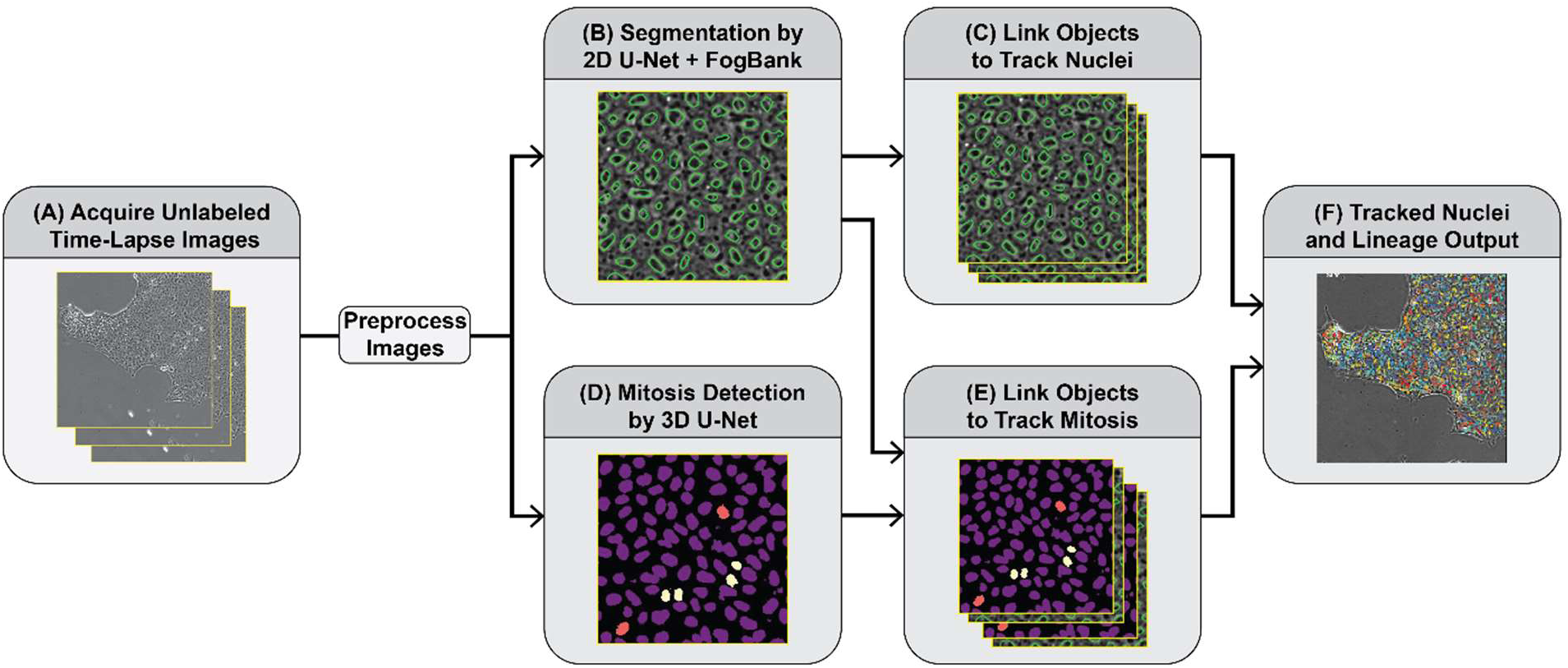
Schematic of the workflow progression for the analysis of time lapse Zernike phase images. (A) Tiled, time-lapse images are acquired and pre-processed by focal plane selection and image stitching. (B) Stitched phase contrast images are inferenced with a 2D U-net, then object separation is performed with the Fogbank algorithm for nuclear segmentation. (C) Nuclear objects are tracked. (D) Time-lapse, stitch phase contrast images are inferenced with a 3D U-Net for mitosis event detection and daughter nuclei identification and detection. (E) Tracked objects from (C) are linked with mitosis events and daughter nuclei from (D). (F) Tracked nuclei and lineages are output. See Methods and Supplemental Material for details.

### Assessing the accuracy of label-free segmentation

The accuracy of inferenced segmentation was calculated after using a linear sum assignment routine with the cost function proportional to both the centroid distances and the overlaps between the segmented GFP reference data and the inferred test set. The distance cutoff between two objects is 15 pixels. The overlap is computed as the intersection over the union of the two objects. The number of false positives or over-segmented objects are normalized by the reference cell count and reported as fraction of additional objects, and the number of false negatives or under-segmented objects are normalized and reported as the fraction of missing objects. We also report F1 scores which were calculated as

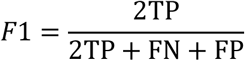

as in (17), where TP is number of true positives, FN is number of false negatives, FP is number of false positives.

### Evaluating the accuracy of automated segmentation for generation of reference data

To estimate the accuracy of automated segmentation of fluorescent nuclei that were used for training and evaluation, we manually annotated centers of nuclei in a small number of fluorescent images. Comparison of automated and manual reference data are discussed in Results and in Supplemental Information Figure 1D and 1E. Errors in object identification were calculated after using a linear sum assignment routine with a cost function proportional to the center distances between the manual annotations and reference fluorescence masks. By comparing manually detected nuclei to nuclei detected with the automated segmentation routine we determined that, while most manually annotated fluorescent nuclei are concordant with automated segmentation results, there were occasionally nuclei that were difficult to detect, where both manual and automated annotation resulted in ambiguous results (Supplemental Figure 1D). By comparing automated reference data with manual data, we determined that the reference data are highly accurate, although some apparent errors occur, mainly due to the difficulty in segmenting overlapping nuclei, resulting in missing objects (Supplemental Figure 1E). The presence of errors in the reference data means that the error rates associated with the inferenced results is slightly overestimated. Nonetheless, the similarity between the reference data and the manually annotated data provides confidence that the reference data produced by automated segmentation can be used to compare and evaluate AI models for label-free segmentation while providing a nearly limitless number of objects that can be sampled.

### Creating training data for label-free mitosis detection

To prepare the training data for the U-Net, GFP data are used to segment cell nuclei for the 2D U-Net with class 0 to designate background and class 1 for nuclei (see *Training the 2D U-Net* section above). Potential mitotic nuclei were then labeled by applying an additional band-pass filter to identify the condensed DNA in the original GFP image followed by a threshold. The pixel values of these objects were designated as class 2. Finally, a tracking routine was used to identify daughter cells. Each object designated as class 2 in frame t_0 is duplicated and then a linear sum assignment routine pairs that object with an object in the subsequent frame t_1. If a cell in t_0 maps to two objects in t_1, then the initial cell is designated as class 2, and the two mapped cells are designated as class 3. A final postprocessing step is applied to designate the object as class 2 (i.e., a mitotic nuclei) in each of the 5 frames before division, and as class 3 (one or two daughter cells) in each of the 3 frames after division. The resulting three-dimensional data were tiled into 128x128x16 images for input into the 3D U-Net.

### Training the 3D U-Net for label-free mitosis detection

To identify mitosis, we built on a previously published 3D U-Net (15) and modified the training data to output a 4-class mask (0 – background, 1 – nuclei, 2 – mitotic nuclei, 3 – daughter cells). For the 3D U-Net, the first two dimensions are the x, y pixels of the image, and the third dimension is consecutive time frames (two-minute separation between frames). The phase contrast images are normalized using z-score normalization, and the final images and reference masks are tiled into 256x256x16 tiles (in the x, y, z directions). During augmentation, a 128x128x16 section of that tile is randomly selected across the x and y directions to match the input of the U-Net. Additional image augmentation was used during the training process in which the input training data were modified by reflections, rotations, and by applying Gaussian blur. The relative weights of the background, non-mitotic nuclei, mitotic nuclei, and daughter cells are 1 to 2 to 20 to 20.Training the 3D U-Net is done in Python using TensorFlow and Keras (https://github.com/usnistgov/WSDOM).

### Post processing of 3D U-Net inferenced results

Mitotic events are detected by combining class 2 and 3 objects as one class– i.e., removing objects that are not mother or daughter nuclei – and counting the number of class 2 and class 3 objects in three dimensions (x, y, time). Potential false positives are filtered by requiring each object to have a volume larger than 300 pixels and persist for at least 10 frames. For subsequent tracking of daughter cells for each event, class 3 designation was used, and the time of mitosis was determined as the time when two objects become identified or when the class 2 object is the largest. If only one class 3 object is identified, then two daughter nuclei are labeled at the major axis positions opposite the mother cell label.

### Evaluation of the accuracy of the 3D U-Net for mitosis detection

Mitoses detected from the 3D U-Net were compared to reference data of 4-class segmentation of GFP fluorescence (see *Training the 3D U-Net* subsection above). To estimate the accuracy of the 4-class automated segmentation results, we manually annotated 629 mitoses across two experiments. The manual data was paired to the 3D U-Net inferenced results using a linear sum assignment routine with the cost function being proportional to the distance between mitosis events in space with an empirically determined spatial cutoff of 15 pixels and a time cutoff of 6 frames.

### Cell tracking with mitosis detection

The inputs for cell tracking are cell nuclei detected using the 2D U-Net, and parent/ daughter positions obtained from the output of the 3D U-Net. Cells are tracked using a tracking routine written in Python (18, 19). The tracking uses a cost function that weights the overlaps as well as centroid distance between objects in consecutive frames to link or map cells across time. In order to reduce the loss of true objects, any unmapped objects are checked for merge events, and any tracked object can remain unmapped for 5 frames after which the track is considered lost. This can result in the appearance of additional segmented objects for single timepoints. Dividing parent/daughter positions are used to track cell lineage and are omitted from the linear sum assignment routine which is applied only to track non-mitotic nuclei.

### Evaluation of cell-tracking workflow

We evaluated our label-free tracking workflow using GFP reference data. To calculate the accuracy, we paired the tracked objects between our reference data and test set (x, y, t, id) in each frame using a centroid-distance cost function as used for the evaluation of the segmentation and mitosis detection. We then calculated the linkage-error rate for each track. A linkage error is when a label-free track id changed, and the reference id did not. The linkage error rate per track is then the sum of these errors divided by the length of the track.

Another intuitive technique to assess tracking accuracy is to compare the long-time measurements between different segmentation models. The metrics that we used are the track lengths (the total number of frames a cell is continuously identified) and the interdivision times (the time between appearance of a daughter cell and its subsequent division), in which a more accurate tracking sequence should have longer track lengths as well as interdivision times that are biologically relevant (i.e., between 9 and 24 hours). We checked the accuracy of the GFP reference data with a small amount of manual annotations.

### Data availability

The data that support the findings of this study are openly available in the NIST Data Repository at https://doi.org/10.18434/mds2-2960. All code and models generated are available at https://github.com/usnistgov/WSDOM.

## Results

The goal of this study is to develop a rapid and robust imaging pipeline that enables in-depth quantitative analysis of unlabeled nuclei in live iPSCs imaged by phase contrast microscopy. Here we develop and evaluate U-Net models that allow segmentation of individual nuclei within iPSC colonies; we also detect and track mitotic cells and the formation of their daughter cells. The output masks of the U-Net models are processed to provide input for tracking and lineage construction of cells for the length of the acquisition.

The general progression of the workflow, as described by the schematic in Figure 1, takes in unlabeled time-lapse transmitted light microscopy images. Two different U-Net operations are applied to the data. The first is inferencing with a 2D U-Net followed by object separation to detect the individual nuclei in each image. The second is inferencing with a 3D U-Net which takes in time-lapse images acquired at 2 min intervals to detect and label mitotic events as well as parent and daughter nuclei. The segmented objects are automatically tracked to link the objects over time and are combined with the labels from the 3D U-Net to generate cell lineages.

Because of the spatial proximity of cells in pluripotent cell colonies, segmenting and tracking individual cells is challenging. Mitosis event detection is also challenging. Adherent mammalian cells in culture frequently round up during division resulting in a high contrast image patterns that can be observed and automatically detected (20). The iPSCs have a more subtle appearance during division, remaining flat with a uniform contrast pattern throughout division. In this work, we have used a mEGFP-HIST1H2BJ cell line to provide fluorescent nuclei that can be segmented and used for training the 2D and 3D U-Nets. High signal-to-noise ratio images can be acquired using this probe allowing for high quality nuclear segmentation with classical image analysis routines. This approach provides the advantage of reducing or eliminating the need for manual segmentation, which in turn allows us to sample a very large volume of reference segmentations for evaluation of the label-free analysis workflow. This latter point is critical because the heterogeneity of cell characteristics and responses demands a large statistical sampling, and this has not typically been accessible particularly in the temporal domain.

The process for development of the 2D U-Net is shown in the schematic in Figure 2A and the steps are described in Methods. Inferenced results were compared to segmented fluorescent images to evaluate the accuracy of the model.

**Figure 2.**
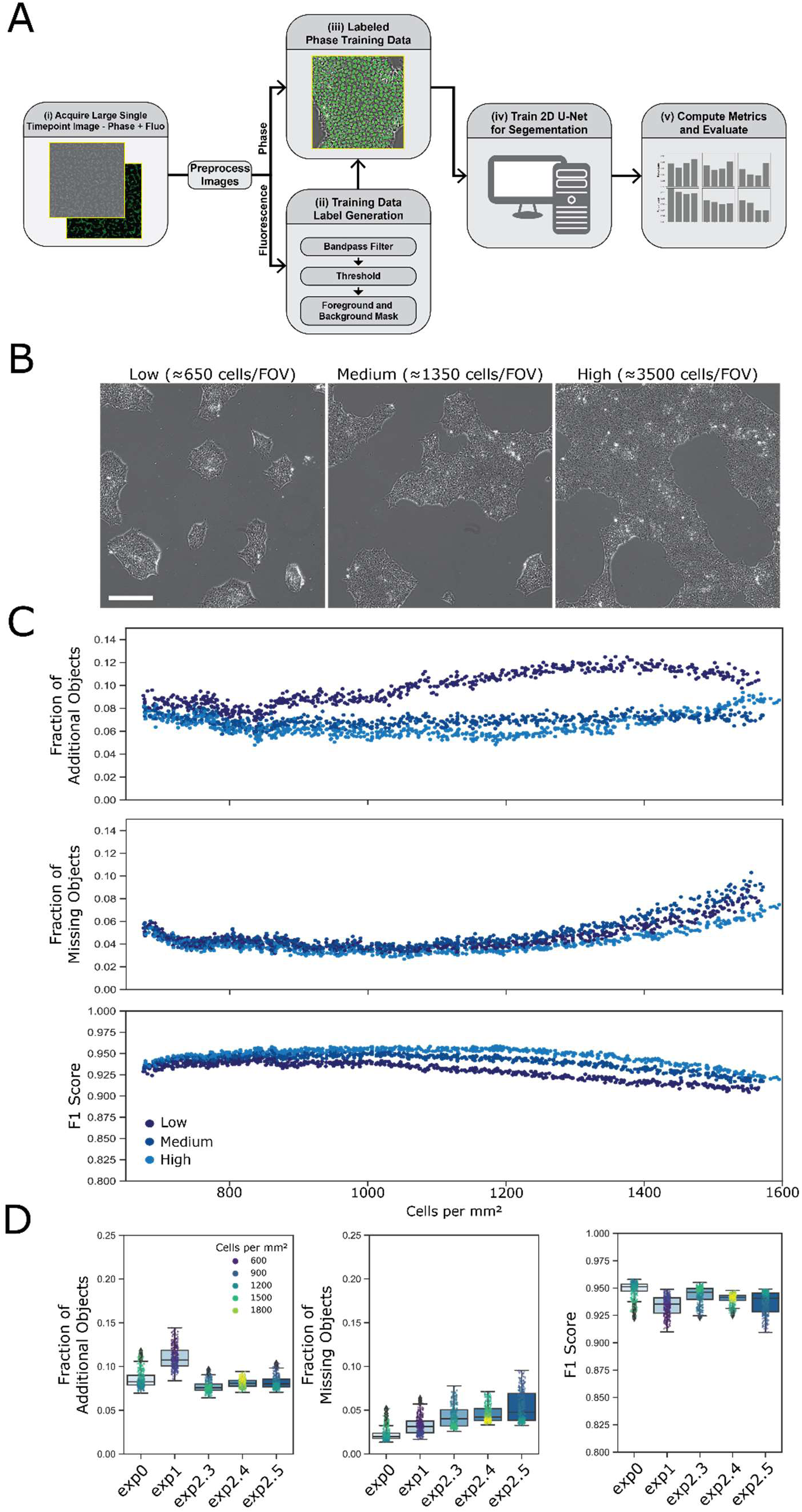
Training and evaluation of 2D U-Nets. (A) Schematic describing the process for model training and evaluation of 2D U-Nets for iPSC nuclear segmentation. (i) A single timepoint image set consists of a phase contrast image created by stitching 100 fields of view (FOV) and a corresponding fluorescence image where nuclei are labeled with H2B-GFP. FOV are acquired and pre-processed to identify the best in-focus z plane and then stitched (see Methods). (ii) The fluorescence images are processed to produce a foreground and background mask. (iii) The processed fluorescence image is combined with the unlabeled image for (iv) training the 2D U-Net. (v) The accuracy of inferencing individual nuclei from phase contrast images is evaluated by comparing the locations of inferenced nuclei with fluorescent nuclei in reference images. (B) Representative phase contrast images of images of cells seeded at relatively low, medium, and high densities. Scalebar shown is 250 µm. (C) The ‘fraction of extra objects’, ‘fraction of missed nuclei’ and the F1 score determined for each frame of one image dataset collected over 20h. Accuracy metrics are plotted versus the measured number of cell nuclei/mm^2^ per frame. The inferenced results were compared for the three models trained on either low (i), medium (ii), or high (iii) cell densities. (D) Using the high-density model, the ‘fraction of extra objects’, ‘fraction of missed nuclei’ and the F1 scores are plotted for five replicate datasets. The first dataset in the plot are the data shown in panel 2C. Each datapoint represents the corresponding error rate for that frame, and the dot color indicates the density of cells in the frame. Tukey box plots indicate summary statistics for each time-lapse dataset.

### Evaluation of 2D U-Nets

#### Comparing models trained with different densities of cells

We created models from three sets of training data containing cells plated at different densities (example images shown in Figure 2B). Depending on the density of cell seeding, time that cells are in culture and natural variability of cultured cells, iPS cells can assume subtle differences in morphologies due to differences in interaction with neighboring cells and adhesion to substrate. To produce a model that is applicable for different datasets and over the times that cells are in culture, we trained models with data from relatively low, medium and high densities of cells. The models were tested against images of cells at different densities to evaluate the generalizability of the models for different cellular contexts. In Figure 2C we show a comparison of the performance of those 3 models with image data that were collected on cells over 20h in culture. The F1 scores, fraction of additional objects and fraction of missing objects were calculated for single frame images taken at different times over 20h of imaging. The data are plotted as a function of the density of cells (number of cells per mm^2^) in the frame. As shown, the models perform similarly over the entire 20h experiment, when cells are at relatively low densities and when cells are at relatively high densities. Comparison of the fraction of additional objects, the fraction of missing objects and the F1 score between 2D U-Net models showed that the performance of the models was not highly sensitive to cell density, and the model trained with the highest-density cells performed nominally better than the others with the ‘fraction of additional objects’ being the largest performance difference between the models. The model generated with a high-density of cells produced an error fraction in this dataset of approximately 0.05 to 0.08 additional objects, and a fraction of approximately 0.03 to 0.06 missing objects, with an F1 score that ranged for 0.92 to 0.95 over the time course of the measurement. The difference in performance of the 3 segmentation models becomes more obvious when cells are tracked over time, as will be discussed later.

#### Comparison of different datasets

We examined different datasets to assess the performance of the high-density model on nominally identical cell preparations. The data in Figure 2D show the accuracy metrics for 5 cell samples from 3 different days, examined over 13h in culture. The data are plotted to indicate the model performance in each frame collected over time and color-coded to indicate the density of cells in the frames. The average F1 score over all 5 datasets is 0.94. These data suggest good concordance in segmentation performance across within-day replicates and experiments on different days. It should be appreciated that many variables are at play when evaluating different datasets in addition to the performance of the U-Net, including unknown biological and experimental variables. The replicability of these data provides confidence that the analysis workflow, including the U-Net, can be robustly applied to nominally different datasets. Segmentation can be compromised by various aspects of the cell culture. These caveats are discussed in Supplemental Material.

To test the robustness of the segmentation and tracking model for different colony-forming cell lines, we applied the model to phase contrast images from the parental WTC-11 iPSC line and the embryonic stem cell line H9. The scores for performance of the model when applied to these 2 cell lines were determined by comparing inferenced results with automated segmentation of reference data created by staining these cells with Spy595-DNA (see Methods). Model performance for these cell lines is similar to what was observed with the H2B-GFP cell line, indicating model generalizability to these other pluripotent stem cell lines (data shown in Supplementary Figure 2).

### Development and evaluation of mitosis detection with a 3D U-Net

A 3D U-Net model was developed to detect mitosis in phase contrast images using training data generated by automated segmentation of the histone H2B-GFP fluorescent nuclei (see schematic in Figure 3A). The result is a 3D U-Net with a four-label output as shown in Figure 3B and Supplemental Video 1 consisting of background (class 0), non-dividing cell nuclei (class 1), nuclei undergoing mitosis (class 2), and daughter nuclei (class 3) immediately after division. The training data for the 3D U-Net were constrained by applying a class 2 label for 5 frames before a division event and applying a class 3 label for 3 frames after the corresponding division event as determined by analysis of the H2B fluorescence data. The output of the 3D U-Net was post-processed to remove spurious mitotic events by filtering for size and duration. Details are provided in Methods.

**Figure 3.**
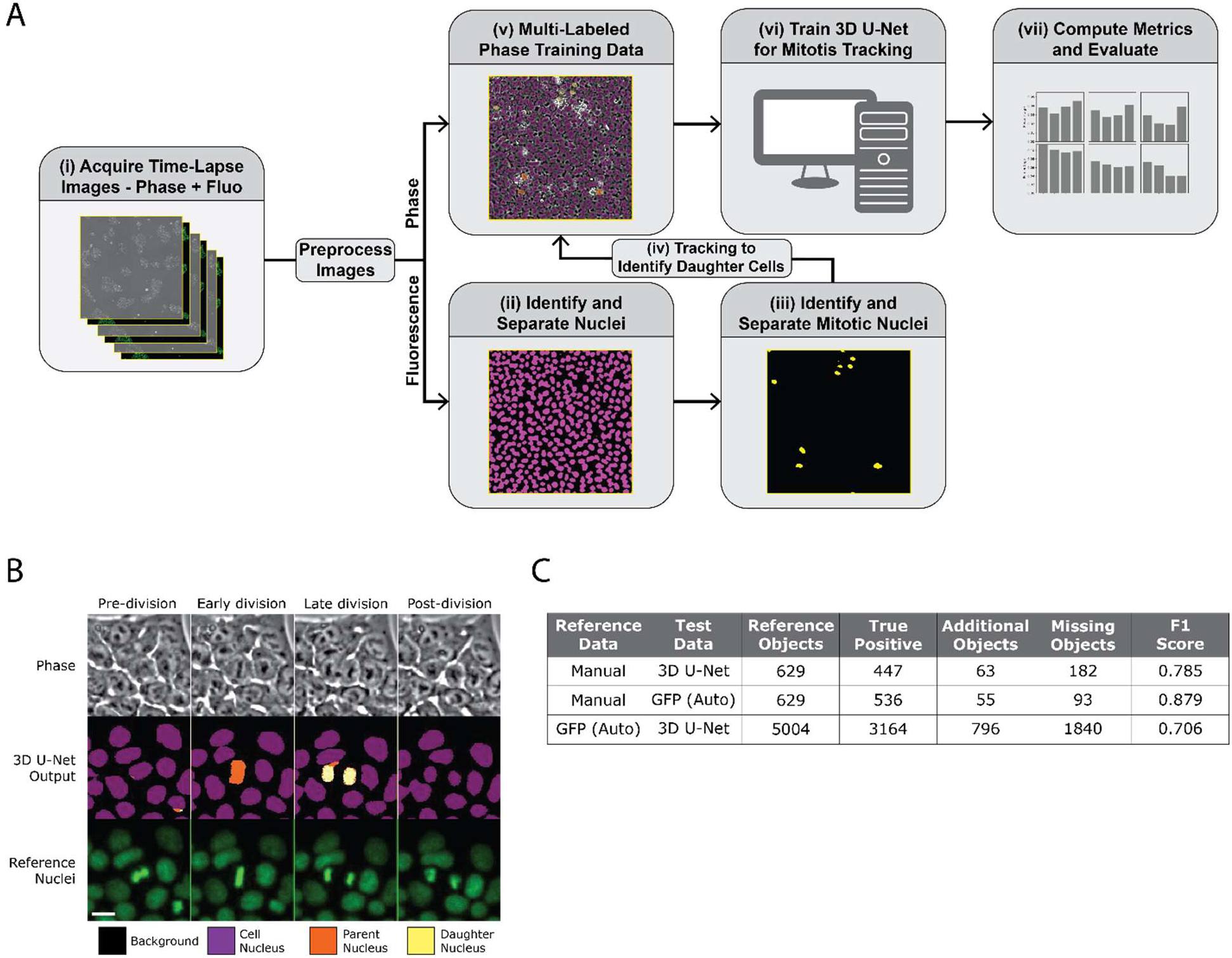
Development and evaluation of mitosis detection with a 3D U-Net. (A) Schematic describing the process for model training and evaluation of a 3D U-Net for iPSC mitosis detection. (i) A time-lapse image set consists of a phase contrast image created by stitching 4 fields of view and a corresponding fluorescence image where nuclei are labeled with H2B-GFP. Images are pre-processed to identify the best in-focus z plane and stitched (see Methods). (ii) The fluorescence images are processed to first identify and classify individual nuclei. (iii) the fluorescence images are processed again to identify and classify mitotic cells. (iv) Afterwards, the different classified cells are tracked over time to identify and classify the daughter cells. (v) The different classes of cells are then combined with the phase contrast images for (vi) training in the 3D U-Net. (vii) The performance is evaluated by comparing the locations of inferenced mitotic events from the phase contrast images with mitotic events identified in H2B fluorescence reference images. (B) Example of a mitotic event occurring in the phase contrast images along with the 3D U-Net classification output with its different class labels and the corresponding fluorescence reference images. Scalebar shown is 15 µm. (C) The calculated error in mitosis detection by the 3D U-Net inferencing when compared to manually annotated reference data, and the error in the fluorescence classification when compared to manually annotated reference data. Comparison of fluorescence reference data (GFP (Auto)) with inferenced (U-Net) data indicates that errors in inferencing are not due to errors in auto classification.

The performance of the 3D U-Net model was directly evaluated by comparing inferenced results with automatically identified mitotic events from fluorescence reference images (Figure 3C). For a dataset consisting of 5004 reference mitotic events that were detected by the automated mitosis detection algorithm applied to the fluorescence data, the 3D U-Net resulted in 3164 true positive mitotic events, 796 additional events, and 1840 missing events for an F1 score of 0.70.

We also examined, with a much smaller amount of manual annotations, the performance of the automated reference data and the inferenced results from the 3D U-Net. Compared with 629 manually annotated mitosis events, the 3D U-Net inferencing resulted in an F1 score of 0.78. Comparing the results of manual annotation with the number of mitosis events detected from the fluorescence data, the F1 score was 0.87. These data indicate that while the fluorescence reference data were in good concordance with manual data, the 3D U-Net tended to miss mitotic events more frequently (182 out of 629 mitotic events were missed by the 3D U-Net). We conclude that the auto-generated reference data is reasonably accurate, and an F1 score for the 3D U-Net of 0.70 is a reasonable estimate. No spatial dependence of the performance of the mitosis detector was observed.

### Label-free tracking of single iPS cells

Time-dependent cell tracking can provide greater insight into evaluation of biological differences between samples and treatments. Additionally, time-dependent cell tracking allows us to filter out spurious short-lived tracks to improve confidence in the data from accurately identified cells. From a biological perspective, the temporal phase contrast image data allow us to access single-cell dynamics in large cell populations over multiple cell doublings. The results of our label-free 2D U-Net segmentation and 3D U-Net mitosis detection routines were used in a cell tracking routine (as described in Methods) in order to follow individual cells and their progeny over time (Supplemental Videos 2 and 3).

Filtering objects that track for only a short period of time (i.e. less than 5 frames or 10 minutes) and removing them from the dataset eliminates spurious objects and other false positive events (Figure 4A). The fraction of additional objects decreases as a function of minimum track length, which indicates that longer tracks are true positive segmentation events, i.e., the objects we are tracking are indeed cells. Filtering short tracks also has implications for cell counting. Removing tracks that are less than 10 minutes in duration results in a cell count that is more consistent with the count from our GFP reference dataset (Figure 4B). Thus, this filtering routine helps to establish greater confidence in both dynamic measurements and also static measurements such as cell count. More stringent filtering criteria can lead to fewer objects tracked but provides greater confidence in the biological implications of the objects’ behavior (Supplemental Figure 3).

**Figure 4.**
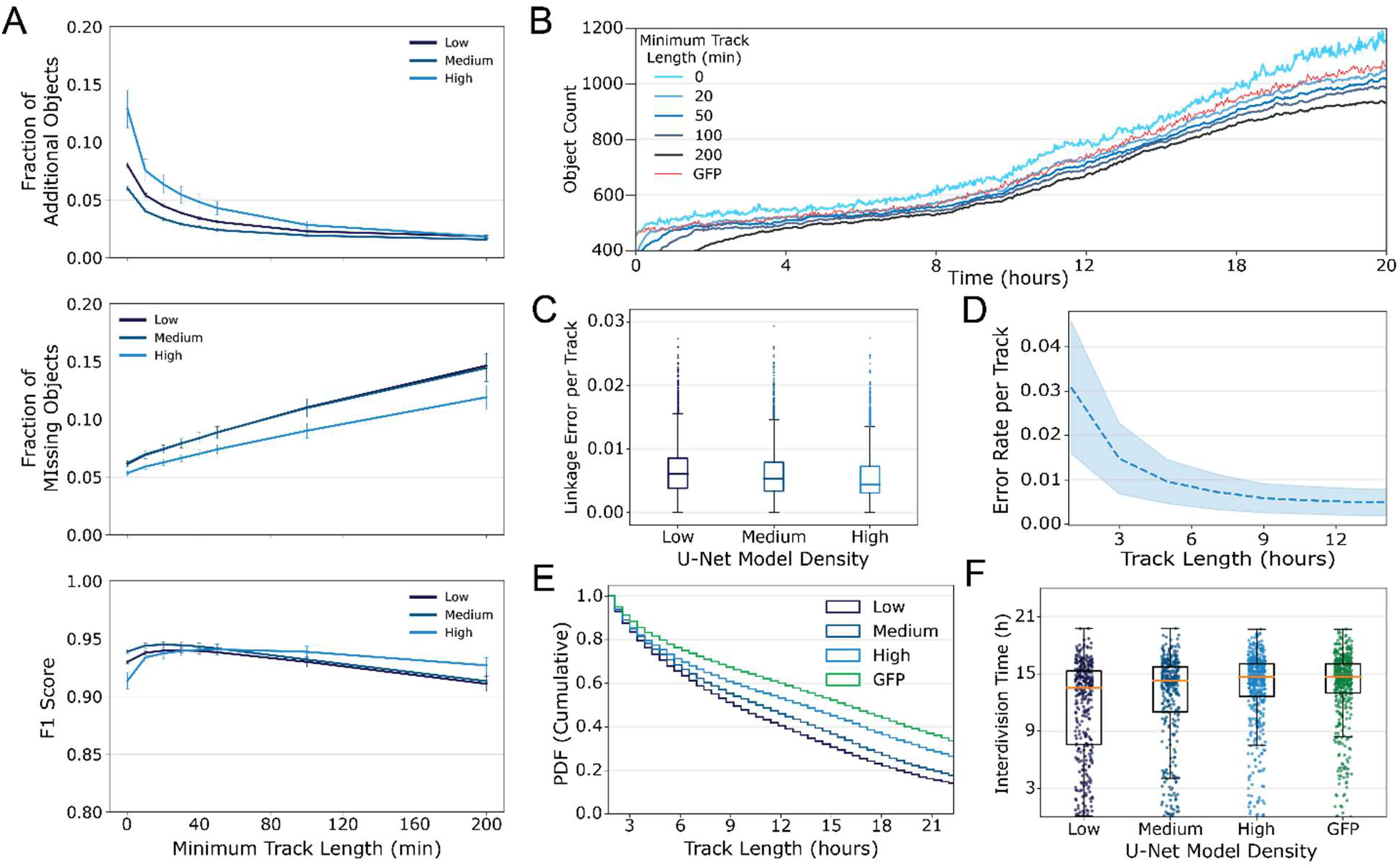
Tracking of segmented iPSC cells. (A) After tracking, eliminating objects with short track lengths reduces segmentation errors. Including only objects that were successfully tracked for times greater than the minimum track length results in datasets with fewer segmentation errors. The model created with the high cell density training data performed better overall than the other models after filtering. (B) Eliminating objects that were tracked for less than 20 consecutive minutes appears to produce a more accurate cell count in inferenced results from the high-density model. The green line is the number of segmented fluorescent cells counted in the corresponding GFP image; that number is nearly identical to the inferred data when a filter of 20 sequential minutes is applied. (C) Tukey boxplot of Linkage Error Rate per track computed by comparing inferenced data from 2D U-Net models trained using either low, medium, or high-density training models and compared with reference tracks created from GFP fluorescence data. Lines in boxes indicate the median error for inferenced tracks was slightly lower from the high-density model. (D) Tracking error rate computed for the high-density 2D U-Net model as a function of track length. (E) Cumulative probability distribution of track lengths inferenced from the different models and from auto segmentation of GFP. (F) interdivision times (hours from a division event to a subsequent division of those daughter cells) for cell tracks computed using the different inferencing models or auto GFP segmentation. Interdivision times that appear to occur within 9 hours are considered physiologically improbable and are likely to be tracking errors. Auto GFP segmentation resulted in 8 of these events compared to 91, 103, and 151 such events for the high, medium and low U-Net models. The numbers of interdivision times that were measured at >9 hours were 441 for the auto GFP segmentation, and 405, 291 and 228 for the high, medium and low U-Net models.

We assessed track lengths as an estimate of the quality of our cell tracking workflow. We quantified cell-tracking accuracy with three different measurements: (1) the linkage error rate per track, (2) the track length distribution and (3) the interdivision times. Using the GFP fluorescence to generate reference tracks, the data in Figure 4C show the linkage error rate computed for an entire dataset (4000 initial cells) to be approximately 0.005 on average, which corresponds to one potential linkage error every 8 hours on average over the course of a single track. Approximately 70% of nuclei within a population can be tracked without linkage errors for more than 8 h. Linkage error rates occur most often in regions of colonies that are difficult to segment (see Supplemental Figure 1). The linkage error rate seems to be lowest for the 2D U-Net model trained on the high-density cell data, which also had shown the best segmentation performance. Using the high-density U-Net data, the linkage error rate as a function of track length is shown in Figure 4D. We observe that tracks longer than 9 hours converge to the error rate of 0.005 – indicating higher confidence in longer tracks. Track length distributions are shown in Fig 4E. Tracking the GFP reference data produces more long tracks than the 2D U-Net-derived inferenced data as expected; however, we see that for the models trained with data from high cell densities, the label-free segmentation results are similar to the GFP results, with the high-density model having the longest tracks. Finally, Figure 4F shows the interdivision times determined from the different models and from the fluorescence data. We make three observations based on these results: (1) the results from the high-density model most closely matches the GFP fluorescence reference dataset, (2) we track more cells from division to division with models developed from data with high cell densities, and (3) the low-density 2D U-Net results in interdivision times less than 9.5 hours – the lower limit of biologically realistic interdivision times (21). This result suggests that using the low-density 2D U-Net leads to more tracking errors than the high-density 2D U-Nets. Accordingly, we use the high-density U-Net for the rest of this study.

### Temporal and spatial analysis of mitosis and migration

Previous studies have demonstrated position-dependent differences between human iPSCs when examining cells at colony edges versus colony centers. Differences in phenotypic characteristics (e.g. focal adhesion attachments) (22–24) and differences in functional responses (e.g. differentiation potential) (25–27) have been observed. These previous studies are limited to single timepoint measurements or averaged responses; the label-free segmentation, mitosis detection and tracking described in Figures 2, 3, and 4 above can be used to quantify spatiotemporal characteristics of many single iPSCs in live cell populations.

The size, shape, and density of colonies vary across the culture substrate. The data in Figure 5 examine whether dynamic single cell characteristics, namely the extent to which cells divide and move, are associated with the location of cells relative to the colony edge. Each colony was segmented according to the distance from the edge of the colony (Figure 5A). As cells grow and divide, each colony increases in size and cells on the interior are increasingly further from the edge. The mean squared displacement (MSD) measured for each individual cell is used to characterize how much a cell moves. There is no clear dependence of cell motion on experiment time, but the motion of individual cells varies spatially within the colonies (Figure 5B). Individual cells in the center of the colony tend to move less than cells near the edge. The mean of the MSDs is plotted as a function of distance from colony edge in Figure 5C and varies from 450 µm^2^ to 140 µm^2^. On average individual cells move the most at the colony edge. Movement decreases as a function of distance from the edge up to 150 µm, when the average movement becomes constant with increasing distance from the edge. This can be observed in the heatmap overlayed on the colonies in Figure 5B. Average mitotic rates do not appear to depend on distance from the colony edge (Figure 5D) and do not correlate with the increased cell motion.

**Figure 5.**
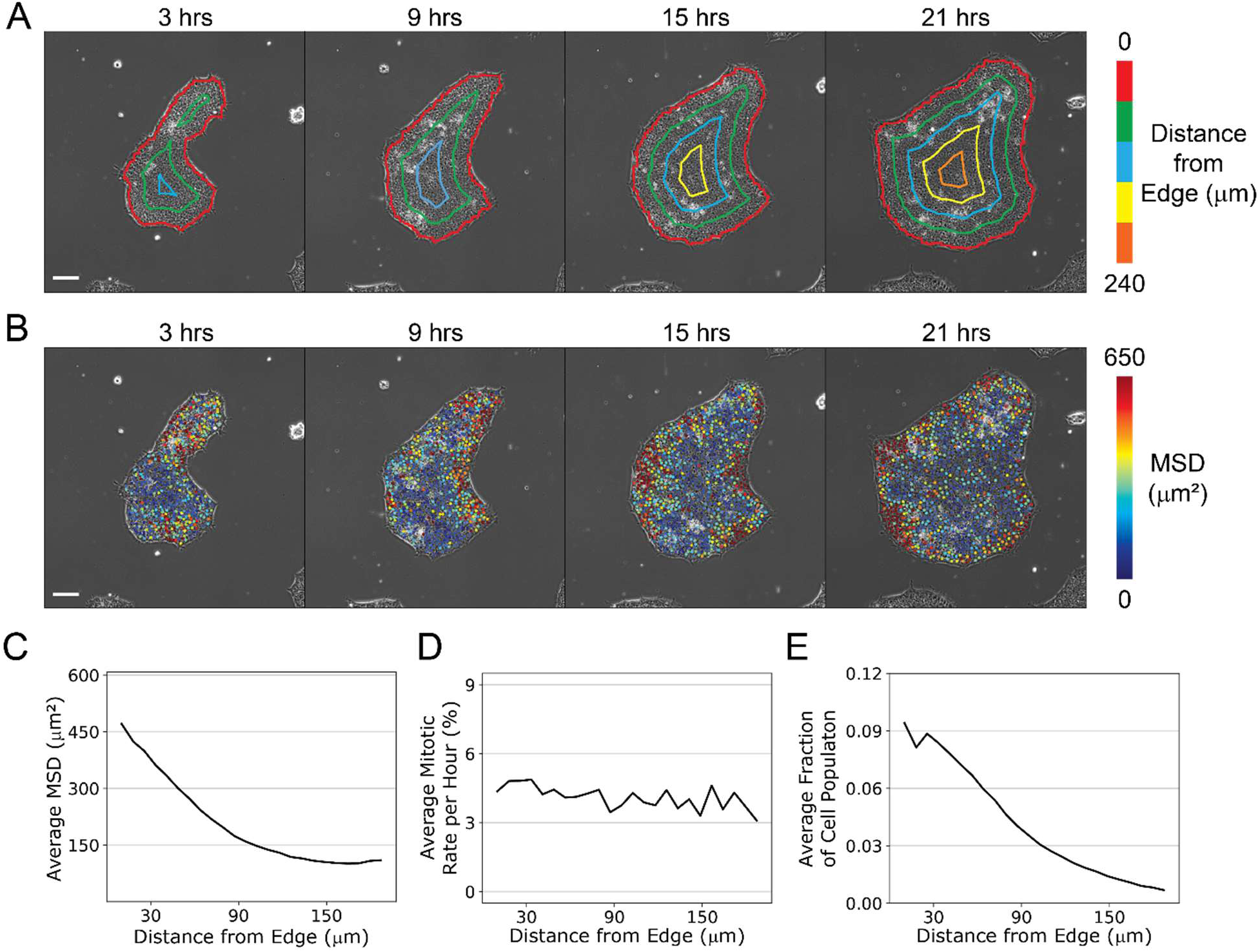
Effect of spatial location within colonies. (A) Phase images of a colony at the times in culture indicated with distances from the edge of the colony labeled with contour lines (scale bar = 100 µm). (B) Images shown in (A) but each cell is colored by its mean square displacement (MSD) calculated over the subsequent hour (scale bar = 100 µm). (C), (D) and (E) show the MSD, mitotic rate and cell fraction as a function of distance from edge of colony, respectively, computed over a time-lapse image series approximately 2200 x 2200 µm over 24 h. Cells move more at the colony edges, but their mitosis rate is unaffected. (Tracks < 100 minutes were not included in this analysis.) No systematic difference in MSD was observed over the time course of the experiment.

### Effect of excitation light exposure on cellular mitosis and cell number

Automated training data for the U-Nets was generated from images of cells with fluorescent nuclei. To confirm that the cells that were exposed to fluorescence excitation light were not significantly different from normal healthy cells, we examined their growth characteristics and compared them to cells that were subjected only to the transmitted light for phase contrast imaging. As shown in Figure 6A, exposure of cells to the minimal intensity of fluorescence excitation light (56 mJ/cm^2^ referred to as 1x) had little effect on cell number over time compared to cells in wells that were only exposed to transmitted light (indicated by the gray line). This result suggests that cells were minimally perturbed by the amount of light they were exposed to at 2-minute intervals over 20 hours. The corresponding data in Figure 6B also indicate that the mitosis rate was apparently unaffected. In contrast, in wells that were exposed to increasing amounts of excitation light, the rate at which cell number increased over time is considerably perturbed. These data are shown in Figure 6A where relative excitation light is indicated in green text as 1.4x, 2.1x, and 3.6x that of the minimal intensity used for generating reference data. Higher intensities of excitation light exposure led to significant cell death that was apparent by manual inspection of images, and by the reduced relative cell numbers as shown by the green lines (example colonies shown in Supplemental Video 4). This was not surprising since radiation is a known inducer of apoptosis (28).

**Figure 6.**
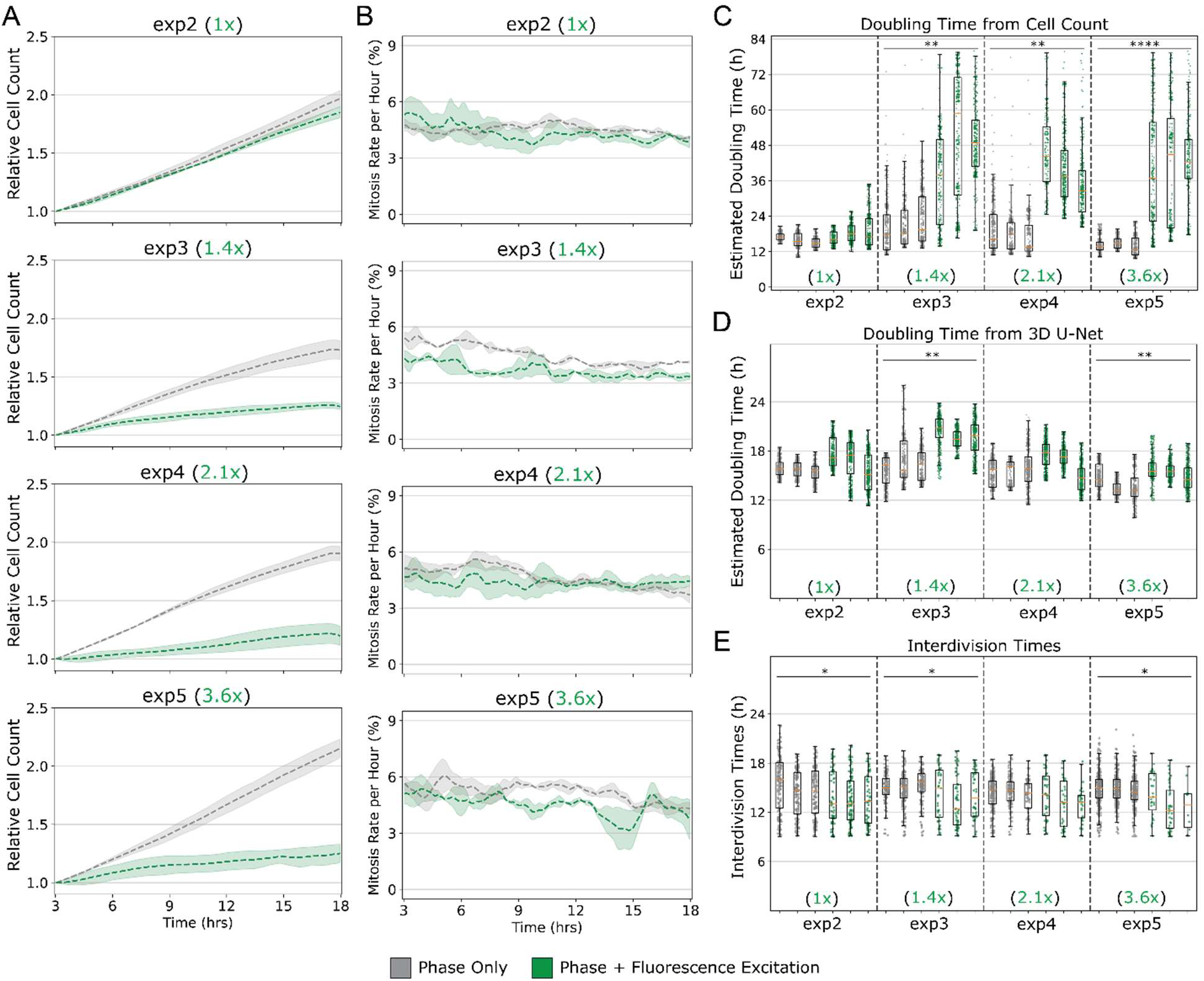
Effect of excitation light exposure on cell division. (A) Cells were exposed only to transmitted light (indicted by grey lines) or to transmitted light and excitation light (indicated by green lines) for 1 sec at 2 min intervals. The relative excitation light is indicated in each panel; 1x is the level of excitation light at which training data were acquired. The number of cells are reported over time relative to the number of starting cells. Shadows around lines reflect standard deviations from replicate samples (n=3). (B) Mitosis rates were determined from the samples analyzed in 6A using the 3D U-Net mitosis detector to determine the total number of mitotic events that occurred within 60 min divided by the initial number of cells in the image in a moving 2-minute time window followed by a 20-frame moving average to reduce noise. (C) Tukey box plots of doubling times calculated from the cell count data corresponding to the relative exposure levels shown in each panel of 6A. Each data point represents a doubling time calculated every two minutes from the numbers of cells in 30 sequential frames (60 min). For the samples exposed to greater than 1x relative flux (green indicators), doubling times are significantly longer than for the other samples. Doubling Time = ln(2)/(ln(N_t_/N_0_)/t). (D) Tukey box plots of doubling times calculated from the numbers of mitotic events determined in samples shown in each panel of Fig 6B corresponding to their relative exposure levels. Doubling Time = ln(2)/(ln(N_t0_+#mitoses)/N_0_)/t). (E) Tukey box plots of interdivision times determined from the samples according to relative light flux exposure. Each dot represents the interdivision time for a single cell. The data associated with higher levels of excitation light exposures resulted in very few cells that progressed from one cell cycle to another, although interdivision times were not different. All data are based on a stitched image of 2x2 FOV with a minimum track length of 100 minutes. All statistical tests were unpaired t-tests using the mean of each replicate comparing within the same experiment, * p < 0.5, ** p < 0.01, *** p < 0.001, **** p < 0.0001.

Interestingly, Figure 6B shows that the number of mitotic cells relative to total cells in the wells was approximately equivalent for wells that were exposed to fluorescence excitation light at any intensity compared to wells that were not exposed to excitation light. The data in Figure 6A show that there are fewer cells in the wells exposed to higher intensity of fluorescence excitation light relative to the untreated wells, suggesting that many cells died as a result of exposure to higher intensities. The smaller number of remaining cells apparently continued to divide at a normal rate as indicated by the data in Figure 6B. This result is similar to studies that have shown that chemical induction of apoptosis results in a wide variability in the extent and rate of cell death within isogenic populations of different cell types (28–30). A comparison of the distribution of division times in cells that were either treated or not treated with an inducer of apoptosis resulted in no significant effect on cell cycle times (28).

The distinction between mitosis rates and cell counts in response to higher intensities of irradiation is evident in Figure 6C and 6D. Figure 6C shows apparent doubling times determined from cell counts. Figure 6C shows that doubling times for cells exposed to the lowest intensity of fluorescence excitation light (shown in the first dataset) are very similar to that of unexposed cells, and the doubling times for unexposed cells over the four different datasets are all similar. However, for the three datasets exposed to higher fluorescence excitation light intensities, the cell counts and apparent doubling times shown in green indicate a strong adverse effect of these higher intensities of light exposure.

It is noteworthy that there is a broad range of doubling time calculated over 60-minute intervals, which is evidence of the variability in susceptibility to the effects of light exposure on apoptosis within the populations. When calculating doubling times based on mitotic events in the remaining cells that were not undergoing apoptosis (Figure 6D), the doubling times are similar to those for unexposed cells, as expected by the small effect of light exposure on measured rates of mitosis, and consistent with previous studies on HeLa cells (28). Manual examination of images confirmed that mitotic cells were accurately identified by the 3D U-Net in these samples. Interdivision times in these populations (Figure 6E) show similar results for all samples, but for samples that were exposed to higher light intensity, many fewer cells are tracked for the length of an interdivision time.

These results are an example of the power of quantitative live cell image data. In the absence of cell-scale data, the low proliferation rate of the population of cells exposed to increasing amounts of light would indicate an apparent phototoxic effect that may be associated with increased death and /or decreased cell division. But the additional mitosis, individual cell distribution, and temporal metrics indicate more nuance in the relationship between mitosis and phototoxicity.

## Discussion

The results of this work allow the use of phase contrast images of iPSCs to provide quantitative cell characteristics, such as mitotic rates, cell migration rates and other dynamic behaviors on a cell-by-cell level and with spatial and temporal resolution. By using phase contrast images for locating and tracking individual cells, fluorescent reporters within cells can be probed intermittently, allowing control over cell exposure to excitation light. Furthermore, the use of phase contrast imaging for tracking cells over their lifetime and the lifetimes of their progeny means that cells can be tracked even if they are not always expressing the reporter. It will be possible to query a very large number of individual cells to study the heterogeneity in temporal responses of cellular features within a population, and the correlations in fluctuations of different features in individual cells. This approach may, for example, provide insight into gene regulatory networks (31). In addition, access to spatial information of individual cells permits the assessment of the effects of changes in culture conditions on cell-cell interactions, the relationship of location within the colony to cell division and motion, and the similarity of timing and location of cell divisions to subsequent divisions of daughter cells.

In this work, we have acquired and analyzed large image datasets, which will be required for uncovering multiparameter correlations in cell populations. Large datasets are also required for the development of reliable models for image analysis, and we demonstrate here the use of fluorescent nuclear probes to automate the process of creating training data. The use of automated segmentation to identify the nuclear foreground from the non-nuclear background allows the creation of very large training and reference data, removing a serious limitation associated with manual annotation. By increasing the number of cells that go into training, we increase the probability that full representation of the heterogeneity of the cell population will be achieved. Similarly, reference data for assessing U-Net model accuracy were also created by automated segmentation of fluorescent images. We have demonstrated that we can, with confidence, use reference data that were generated with classical algorithms to compare models. The 2D U-Net for nuclear segmentation was trained from single time point images, a process that can be easily replicated if needed. The 3D U-Net which identifies classes of mitosing cells requires time-lapse images of sufficient frequency; more training data would be required to achieve the same segmentation accuracy from the 3D U-Net as the 2-D U-Net.

We have demonstrated some examples of the sensitivity of parameters used in preparing the data and selecting the models to the inferenced outcome, and some of the caveats are discussed in Supplemental Material. We have clearly not examined all the possible parameters that could be explored. Complicating the analysis process is the variability that is typical in biological samples from one preparation to another. Despite attempts to be consistent, clearly there are subtle unknown differences in cell and material handling that result in different culture behaviors, particularly from one day to another. These variations may be convoluted with operation of the models, and occasionally there are culture conditions that preclude satisfactory application of any model, such as floating debris that obscures underlying cells. In addition, one might want to create filters to recognize bad frames where, for example, cells were occluded by floating debris. For example, we found that applying temporal filters to remove objects that are tracked only for short times is a strategy for examining only those cell objects that are tracked with a very high level of confidence.

Ultimately, the use of large volumes of reliable quantitative real time images will aid not only cell biology research, but also the monitoring and evaluation of cell manufacturing processes. Especially for manufacturing applications, image analysis must be rapid, automated, and demonstrated to be reliable. Generalizability is also important for adapting the model to different applications. While we used a GFP-producing reporter cell line for training and reference data, one could use different nuclear (or other) markers for model development. We have not examined generalizability exhaustively in this study, but we did observe that the 2D U-Nets performed well on other cell lines, namely the parental WTC-11 cell line and a human embryonic stem cell, H9, using a different fluorescent nuclear probe for reference.

## Conclusion

In this work we have established that single cells in iPSC colonies can be identified and tracked for long times with good accuracy, and that automated segmentation and classification of nuclei based on fluorescent probes can obviate the need for manual annotation. The use of automated annotation in combination with high-speed acquisition rates will enable the efficient development of multiple models to analyze cells in different biological states and conditions over time. This is important for creating useful AI models with high accuracy, and it allows the sampling of many cells as required to capture cell-to-cell variability. This capability will provide insight into complex biological processes by enabling the quantitation of correlations between many features of iPSC systems that are accessible by real-time imaging, and lead to better understanding of the controlling factors of emergent cellular properties and how to predict and direct them. In addition to the cellular characteristics, we have examined here (motility, division rates and interdivision times), this work will facilitate the monitoring of many metrics of interest over space and time, including intracellular structural features, expression of transcription factors and other markers of gene expression, and markers of functional response such as differentiation.

## Acknowledgements

Disclaimer: Commercial products are identified in this document in order to specify the experimental procedure adequately. Such identification is not intended to imply recommendation or endorsement by the National Institute of Standards and Technology, nor is it intended to imply that the products identified are necessarily the best available for the purpose.

