## Supplemental Material for "High-volume, label-free imaging for quantifying single-cell dynamics in induced pluripotent stem cell colonies"

CONTENTS

1. *Supplemental text*
    a. *Image acquisition (pg. 2-3)*
        i. *Microscope System*
        ii. *General image acquisition workflow*
        iii. *Single time point imaging for collecting image data for training 2D U-Nets*
        iv. *Live cell imaging*
    b. *Imaging instrument benchmarking (pg. 3)*
    c. *Image pre-processing (pg. 3)*
    d. *Considerations for evaluating confidence in AI results (pg. 4)*
    e. *Mean square displacement analysis (pg. 4)*
2. *Supplemental movie captions (pg. 5)*
3. *Supplemental figures (pg. 6-9)*
4. *References (pg. 10)*

**Disclaimer**: Commercial products are identified in this document in order to specify the experimental procedure adequately. Such identification is not intended to imply recommendation or endorsement by the National Institute of Standards and Technology, nor is it intended to imply that the products identified are necessarily the best available for the purpose.

### **Image Acquisition**

#### Microscope system

Images were collected with a Zeiss Axio Observer.Z1 microscope (431007-9901-000, Carl Zeiss USA, Thornwood, NY) equipped with motorized x-y stage (MLS203-2, Thorlabs, Newton, NJ) and an ORCA-Fusion BT Digital CMOS camera (C15440-20UP, Hamamatsu, Japan) for image capture. Samples were imaged with phase-contrast illumination using a 590nm LED source (M590L4-C4, Thorlabs, Newton, NJ) set to Kohler conditions corresponding to the Zeiss 10X, 0.3NA objective (420341-9911-000, Carl Zeiss USA) used for imaging. Fluorescence excitation of samples was done using an LED excitation source (pE-4000, CoolLED, Andover, UK) through filter set 38 HE (489038-9901-000, Carl Zeiss USA) for mEGFP imaging and filter set 43 HE (489043-9901-000, Carl Zeiss USA) for Spy595-DNA imaging. A spatial calibration target was used to determine that each pixel is equivalent to an area of 0.401 $\mu m^2$. During automated acquisition, the microscope hardware and x-y stage were controlled with Inscoper software (Inscoper, Rennes, France). Cell experiments were performed with controlled temperature and $CO_2$ using a microscope incubation chamber (XLmulti S1 DARK LS, PeCon GmBH, Erbach, Germany).

#### General image acquisition workflow

The general automated acquisition protocol was as follows: 1) A plate focus map was created by scanning a random sample of positions across the imaging area, autofocusing on each position, then estimating the correct focus position for each field of view. 2) The stage moves from field to field to the z-position determined from the plate focus map. Each field overlaps adjacent fields by 10% to facilitate stitching of all fields into a single composite image. At each field, 7 images are acquired above and below the predicted focus position with a z-spacing of 2.5 μm for phase-contrast imaging and a single plane at the predicted focus position for fluorescence.

#### Single time point imaging for collecting image data for training 2D U-Nets

In this study, we used microscopy instrumentation control that permitted rapid data acquisition, which allowed us to efficiently collect large amounts of image data with a 10x objective for training U-Nets for identification of individual iPSC nuclei. We trained 2D U-Nets with 230,000,000 pixels of image data, which translated up to 215,000 cell/nucleus objects. Approximately 0.2 hours of imaging were required to collect the cell data used for training this 2D U-Net, compared to approximately 6 hours of imaging required for our previous study (1). We automated the annotation of these large training data sets using traditional algorithms to segment fluorescent nuclei, which provided much more training data than would have been available by manual annotation.

Live cell imaging

For live cell experiments, paired phase-only and phase + fluorescence excitation imaging was done for each condition with triplicate wells. Exposure times were kept consistent between experiments with phase-contrast imaging at 100ms and fluorescence excitation at 1000ms. Fluorescence excitation intensity was varied for the different dosages. The imaging sequence involved acquiring a 2x2 tile in each well of a 6-well plate in phase-contrast or both phase-contrast and fluorescence every 2 minutes for 20+ hours. This results in a minimum of 100,000 images to be processed and analyzed per well. All data is written locally before being actively transferred to a larger network attached storage for downstream processing and analysis.

**Imaging instrument benchmarking**

Periodically (approximately once per month), we performed a series of benchmarking measurements to evaluate the operation of the microscopy system. Incident illumination power at the sample level was measured using a digital optical power meter and slide photodiode power sensor (PM100D and S170C, Thorlabs, Newton, NJ). Testing of stage functionality and repeatability, field distortion, field uniformity, lateral co-registration, and z-stack drift was performed by imaging with the Argo-LM slide and analyzed using the Daybook 3 software (Argolight, Pessac, France). Optical spatial calibration was completed using a cross micrometer (60210-13PG, Electron Microscopy Sciences, Hatfield, PA). A positive 1951 USAF resolution test target (R3L3S1P, Thorlabs) was used to confirm optical resolution. Camera alignment with the optical system was completed using a line grid target (62-536, Edmund Optics, Barrington, NJ).

**Image pre-processing**

For each time point, the microscope takes multiple field of view images (2000x2000 pixels) over a large, multi-field of view area with 10% overlap between fields of view (also referred to as tiles). Each field of view contains 7 phase-contrast images at varying z planes and a single fluorescence image. Before stitching, the most-in-focus phase image is obtained by selecting the sharpest image, as determined as the z-position with the highest average image derivative computed using Roberts operator. Then, the phase-contrast images are stitched using MIST(2). Fluorescence images are stitched using the same stitching vectors obtained from the phase-contrast channel. Stitching vectors are smoothed across time to avoid jumps that could be due to artefacts like floating debris.

**Considerations for evaluating confidence in AI results (i.e. caveats)**

Biological samples can be variable and future work could consider filters that would flag and/or remove data that are not likely to be accurately segmented or tracked because of the following features :

- Retraction of small colonies as a result of media change which creates areas of cells that are temporarily bright, rounded, and difficult to segment. As a result, some earlier frames in a time series can return segmentation results that are less accurate, but later time points were well analyzed.
- Debris in the FOV: jumps in tracking that could be due to artifacts like floating debris. We employed stitching vectors that were smoothed across time to avoid this. In some cases where there is significant cell death, inferenced segmentation is impossible because of obscuration of cells in the phase contrast images.

**Mean square displacement analysis**

The average mean squared displacements (MSD) reported in Figure 4C were calculated as $MSD(d) = \langle [p(t+\Delta t) - p(t)]^2 \rangle_d$, where $p$ is x,y position of the centroid of a detected nuclues, $\Delta t = 1\ h$ is the time lag applied for determining the displacement, $d$ is the distance from the edge of the colony, and the bracket denotes an average over all $d$. An MSD is determined at all possible $\Delta t = 1\ h$ increments for each cell, and then, MSD values are binned according to distance from the colony edge as indicated in Fig. 4C. We assume the MSD does not depend on time, so we can average over 1 h increments for each cell across the experiment. $\Delta t = 1\ h$ was chosen to reduce the contribution of measurement uncertainty associated with localizing the nuclear object centroid to a single x, y coordinate. In addition to the analysis presented in Figure 4, MSD values were examined for trends as a function of experiment duration and no trend was found. MSD analysis is reported for all tracked nuclear objects from image data in exp0.

**Supplemental video captions:**

**Supplemental Video 1.** Timelapse images showing representative output of the 3D U-Net. The field of view shown is the same as shown in Supplemental Movie 3 for comparison. The class outputs are as follows: nucleus = purple, mitotic nucleus = orange, daughter nucleus = yellow, and background = black. Time is shown as h:min.

**Supplemental Video 2.** Time-lapse images of exp0 showing the phase contrast channel (greyscale) and overlayed with the inferenced and tracked nuclei (color). Individual tracked cells are identified as objects that maintain the same color from one frame to the next. The initial and final cell counts were 3983 and 11358, respectively. Debris on the detector can be seen to create occasional spurious nuclear objects by the 2D UNET; detecting such imperfections in the imaging system and correcting them is important for minimizing inaccuracies. Upon removal, the generation of spurious objects due to the debris was eliminated. Time is shown as h:min.

**Supplemental Video 3.** Sub-region of Supplemental Movie 1 shown with higher spatial resolution. The phase contrast channel (greyscale) is overlayed with the inferenced and tracked nuclei (color). Individual tracked cells are identified as objects that maintain the same color from one frame to the next. The movie begins with two colonies in the center of the field of view. Time is shown as h:min.

**Supplemental Video 4.** Selected timelapse phase contrast images of iPS cells exposed to varying levels of fluorescence excitation. From right to left: 0x, 1x, 1.4x, 2.1x and 3.6x light dose. Time is shown as h:min.

### Supplemental Figures

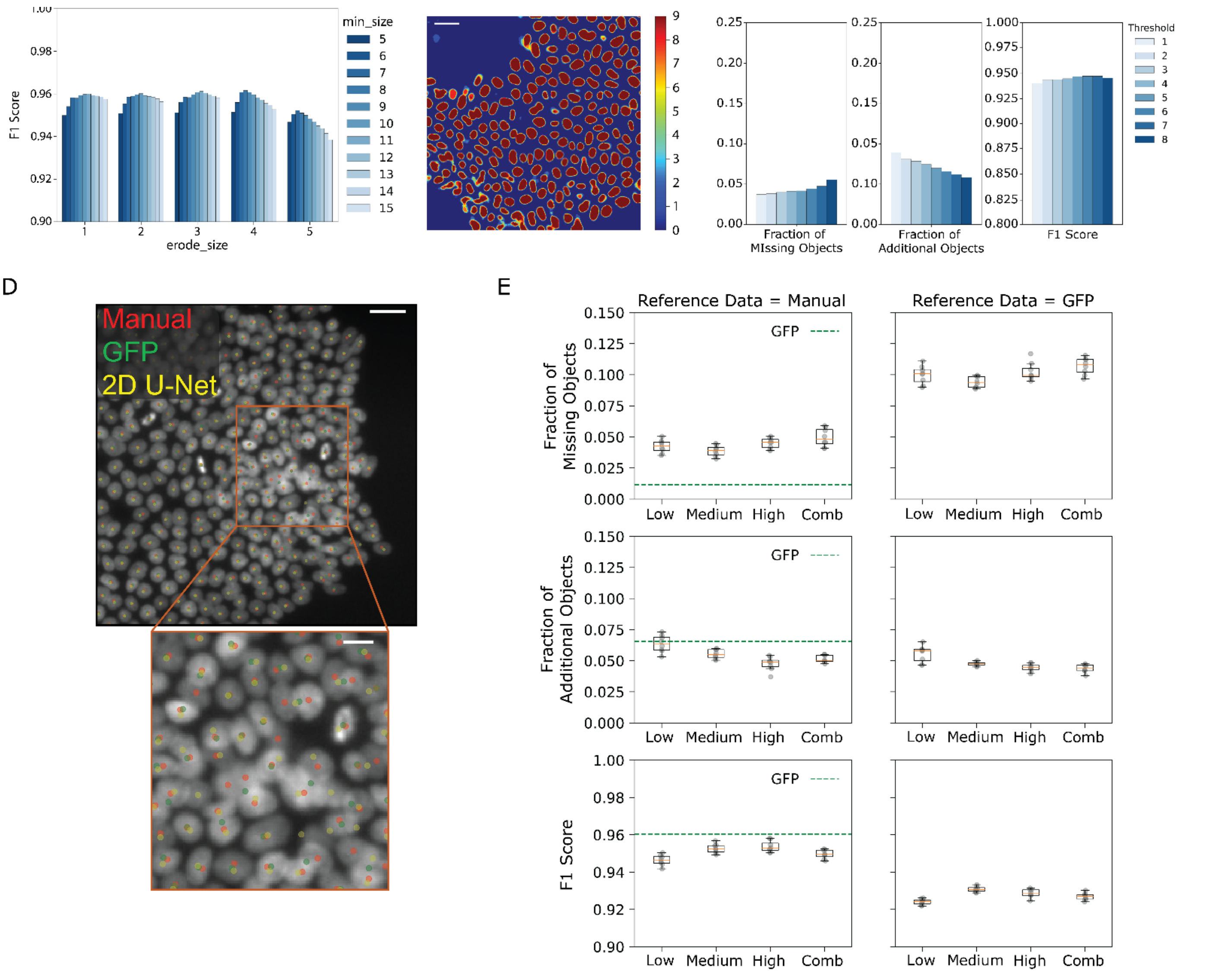


**Supplementary Figure 1. (A)** The effect of relevant Fogbank algorithm parameters on the F1 score are evaluated. The Fogbank algorithm is applied after the 2D U-Net to separate two or more nuclei that share a boundary and are considered one object. The 'erode_size' parameter is varied from 1 to 5 and for each 'erode_size' value, the 'min_size' parameter is varied from 5 to 15. The highest F1 scores for segmentation accuracy can be obtained with 'erode-size' in the range of 1 to 4 and 'min_size' in the range of 8 to 10. **(B)** Representative sub-image from exp0 in Figure 2 panel D showing variability associated with training and inferring with the 2D U-Net is shown (scale bar = 25 µm). The color scale indicates the number of times a trained U-Net inferred that a pixel was classified as a nucleus. Many pixels exhibit high model concordance (9/9), while other pixels exhibit larger discordance. **(C)** A threshold was applied to the image data in exp0 (a representative sub-image shown in **(B)**) and the model performance scores: 'Fraction of Missing Objects', 'Fraction of Additional Objects' and 'F1 Score' are plotted as a function of the threshold value. As expected, the 'Fraction of Missing Objects' increases with threshold value, the 'Fraction of Additional Objects' decreases with threshold value, and the ‘F1 Score’ is highest for intermediate threshold values. **(D)** Three sets of detected nuclei data are shown: Nuclei detected by manual inspection of GFP fluorescence images (red dots), nuclei detected by classical image analysis of GFP fluorescence images (green dots) and nuclei detected by AI-based analysis of the phase contrast images (yellow dots). Scale bar = 10 µm. Many image regions illustrated high concordance between the three datasets, whereas the inset square highlighted in orange illustrates a region of high discordance (scale bar = 10 µm). The GFP fluorescence-based automated image analysis tends to merge nuclear objects compared to the manual annotations and the AI-based analysis of the phase contrast images. **(E)** The model performance scores: ‘Fraction of Missing Objects’, ‘Fraction of Additional Objects’ and ‘F1 Score’ are plotted for inferenced data from 4 different U-Net models using low, medium, high and combined data. The reference data for computing the scores was derived from either nuclei detected by inspection of the GFP fluorescence images or the nuclei detected by classical image analysis of the GFP fluorescence images. The relative performance of the models is similar using either approach for generating reference nuclei.

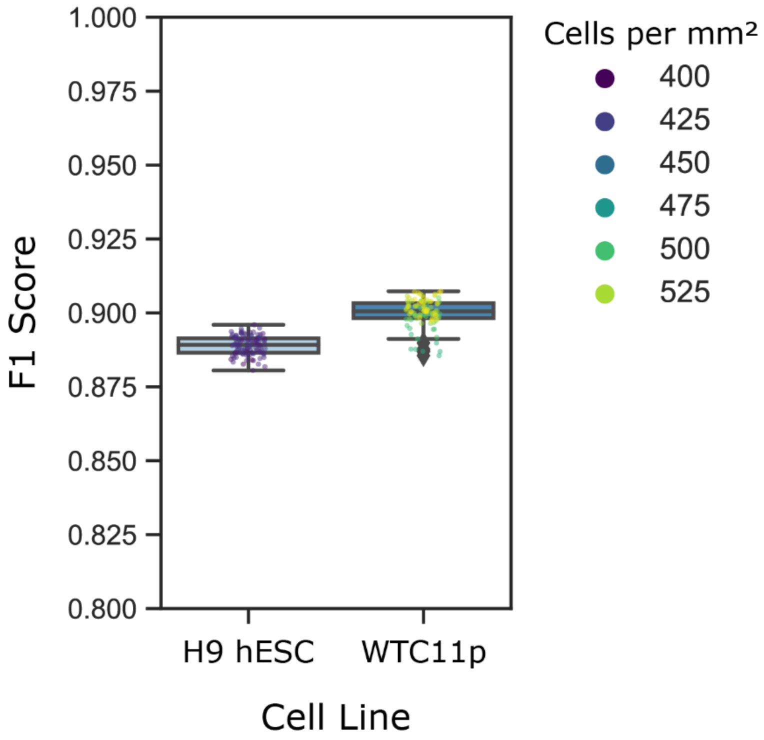


**Supplementary Figure 2.** Using the high density model, the F1 scores are plotted for both H9 hESC (exp6.0) and the parental WTC11 lines (exp6.1). Each datapoint represents the corresponding error rate for that frame, and the dot color indicates the density of cells in the frame. Tukey box plots indicate summary statistics for each timelapse dataset.

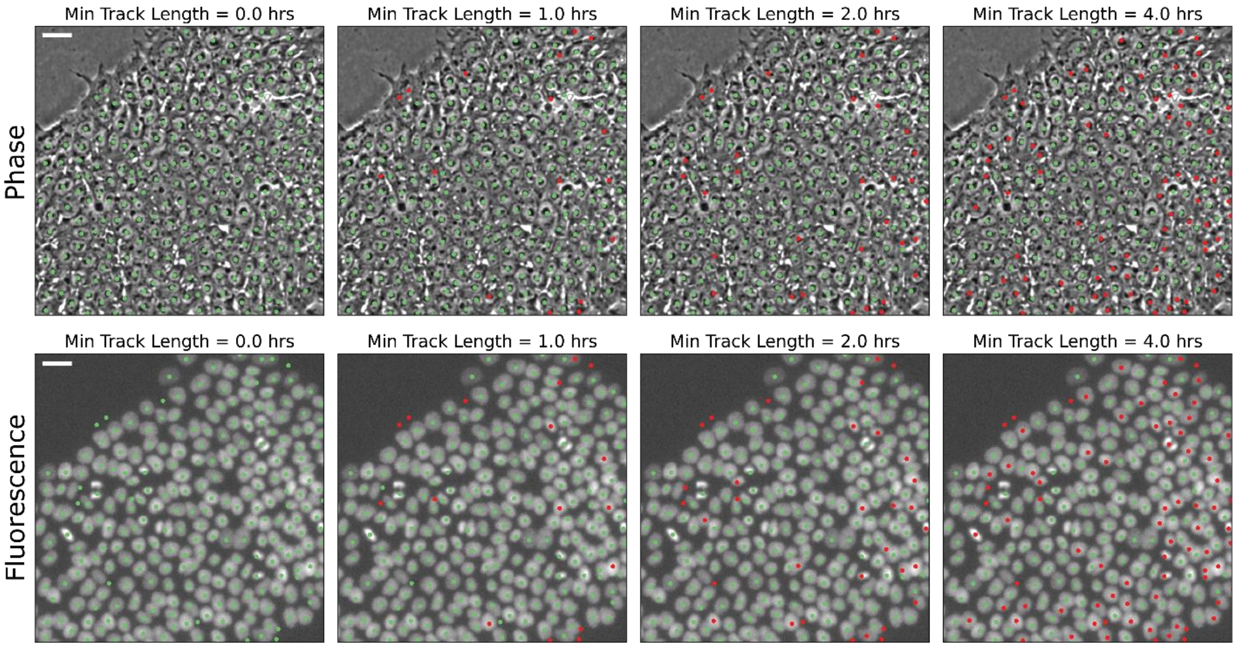


**Supplementary Figure 3.** Example of track filtering based on minimal track length. Green dots represent tracks that have a track length greater than the minimum track length time and are kept. Red dots represent tracks that are filtered out due to having a track length smaller than the minimum track length time. Corresponding phase and fluorescence images are shown for minimum track length times of 0, 1, 2, and 4 hours. Scale bars = 25 µm.
